# The E3 ubiquitin ligase MGRN1 targets melanocortin receptors MC1R and MC4R via interactions with transmembrane adapters

**DOI:** 10.1101/2025.03.25.645338

**Authors:** Pragya Parashara, Lei Gao, Alyssa Riglos, Sonia B. Sidhu, Dorothy Lartey, Tessa Marks, Carys Williams, Grace Siauw, Anna I. L. Ostrem, Christian Siebold, Maia Kinnebrew, Michael Riffle, Teresa M. Gunn, Jennifer H. Kong

**Affiliations:** Department of Biochemistry, School of Medicine, University of Washington, Seattle, Washington, USA; Division of Structural Biology, Wellcome Centre for Human Genetics, Nuffield Department of Medicine, University of Oxford, Oxford, OX3 7BN, UK; Department of Biochemistry, Stanford University School of Medicine, Stanford, California, USA; Department of Genome Sciences, School of Medicine, University of Washington, Seattle, Washington, USA; McLaughlin Research Institute and Touro University College of Osteopathic Medicine, Great Falls, Montana, USA

## Abstract

E3 ubiquitin ligases play a crucial role in modulating receptor stability and signaling at the cell surface, yet the mechanisms governing their substrate specificity remain incompletely understood. Mahogunin Ring Finger 1 (MGRN1) is a membrane-tethered E3 ligase that fine-tunes signaling sensitivity by targeting surface receptors for ubiquitination and degradation. Unlike cytosolic E3 ligases, membrane-tethered E3s require transmembrane adapters to selectively recognize and regulate surface receptors, yet few such ligases have been studied in detail. While MGRN1 is known to regulate the receptor Smoothened (SMO) within the Hedgehog pathway through its interaction with the transmembrane adapter Multiple Epidermal Growth Factor-like 8 (MEGF8), the broader scope of its regulatory network has been speculative. Here, we identify Attractin (ATRN) and Attractin-like 1 (ATRNL1) as additional transmembrane adapters that recruit MGRN1 and regulate cell surface receptor turnover. Through co-immunoprecipitation, we show that ATRN and ATRNL1 likely interact with the RING domain of MGRN1. Functional assays reveal that MGRN1 requires these transmembrane adapters to ubiquitinate and degrade the melanocortin receptors MC1R and MC4R, in a process analogous to its regulation of SMO. Loss of MGRN1 leads to increased surface and ciliary localization of MC4R in fibroblasts and elevated MC1R levels in melanocytes, with the latter resulting in enhanced eumelanin production. These findings expand the repertoire of MGRN1-regulated receptors and provide new insight into a shared mechanism by which membrane-tethered E3 ligases utilize transmembrane adapters to dictate substrate receptor specificity. By elucidating how MGRN1 selectively engages with surface receptors, this work establishes a broader framework for understanding how this unique class of E3 ligases fine-tunes receptor homeostasis and signaling output.

## INTRODUCTION

Mahogunin Ring Finger 1 (MGRN1) is a multifunctional E3 ubiquitin ligase involved in diverse cellular processes, including mitochondrial quality control, endosomal trafficking, and the regulation of signaling pathways. In mouse models, loss of function mutations in *Mgrn1* lead to severe phenotypes, including abnormal pigmentation, birth defects, neurodegeneration, and metabolic dysfunction, underscoring its critical role in maintaining cellular homeostasis (**Table 1**). Despite these findings, little is known about how this versatile E3 ligase identifies and targets specific substrates for ubiquitination. One area of particular interest is how MGRN1 regulates surface receptor abundance. Although the precise mechanism remains incompletely understood, we know that MGRN1 and its vertebrate-specific paralog RNF157 represent a novel pair of membrane-tethered E3 ligases that interact with surface-bound adapters to selectively ubiquitinate receptors within the plasma membrane. We previously demonstrated that MGRN1 binds to the cytoplasmic tail of the single-pass transmembrane adapter Multiple Epidermal Growth Factor-like 8 (MEGF8) to promote the degradation of the G-protein coupled receptor (GPCR) Smoothened (SMO) within the Hedgehog signaling pathway^1,2^. However, the wide range of phenotypes observed in *Mgrn1* deficient mice extends beyond defects in Hedgehog signaling, suggesting that MGRN1 may also regulate other surface receptors mediated through interactions with additional transmembrane adapters.

**Table 1:**
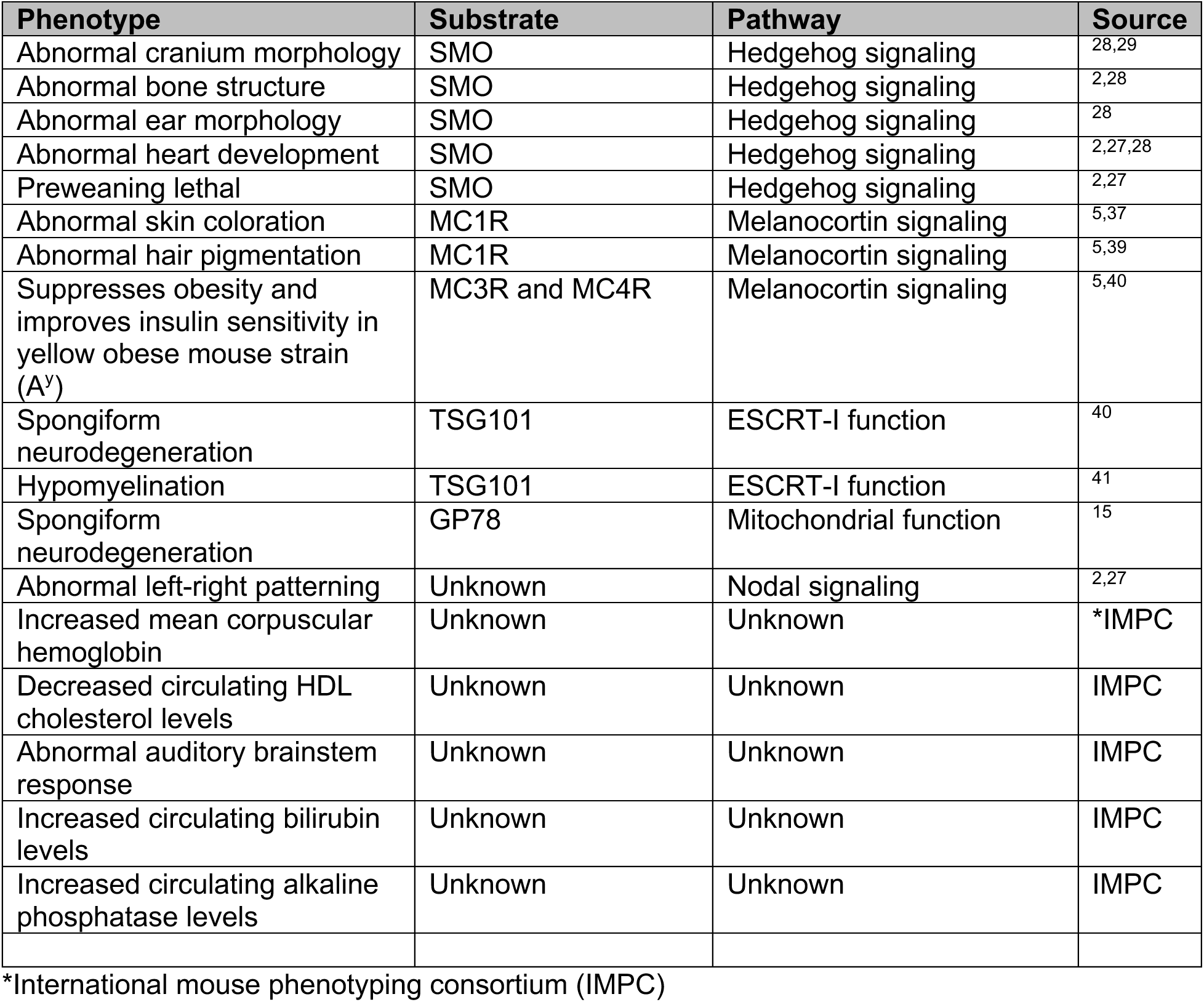
Table of phenotypes in *Mgrn1* mutant mice.

The melanocortin receptor (MCR) family is comprised of five GPCRs: MC1R, MC2R, MC3R, MC4R, and MC5R, which are expressed in different tissues. These receptors regulate a broad spectrum of physiological processes, including pigmentation, stress response, energy homeostasis, and appetite control. Epistasis experiments in mice have shown that MGRN1 inhibits MC1R and MC4R signaling^3–5^ (**Supplementary Figure 1A** and 1B), with prior studies reporting interactions between MGRN1 and MC1R, MC2R, and MC4R^6,7^. However, conflicting data have left the mechanisms underlying this inhibition unclear. Studies in mice suggest that MGRN1 regulates MC1R through its E3 ubiquitin ligase activity^8^. In contrast, other studies propose that MGRN1 modulates MCR function independently of ubiquitination, possibly by competing with G⍺s proteins for receptor binding^6^. These discrepancies underscore the need to clarify how MGRN1 modulates MCR signaling.

To address this gap, we provide evidence that MGRN1 interacts with transmembrane adapters to modulate the surface abundance of MC1R and MC4R via ubiquitination, utilizing a mechanism similar to its regulation of SMO^2^. Using mass spectrometry and biochemical assays, we demonstrate that MGRN1 binds directly to the transmembrane adapters Attractin (ATRN) and Attractin-like 1 (ATRNL1). Through ubiquitination assays, we show that MGRN1 requires interaction with either ATRN or ATRNL1 to ubiquitinate and degrade MC1R and MC4R. Furthermore, we show that the loss of *Mgrn1* results in elevated surface and ciliary MC4R levels in NIH/3T3 fibroblast cells. Similarly, in melanocytes, a loss of MGRN1 leads to increased surface MC1R and enhanced signaling activity, as measured by elevated eumelanin production. Collectively, these findings support a new model in which MGRN1 functions as a membrane-tethered E3 ligase to regulate surface GPCR abundance and fine-tune signaling sensitivity in receiving cells.

## RESULTS

### MGRN1 interacts with ATRN and ATRNL1

MGRN1 and its vertebrate-specific paralog RNF157 are among a few known E3 ubiquitin ligases that interact with membrane-bound adapters to modulate the abundance of surface-bound signaling receptors^9^. MGRN1 and RNF157 exhibit functional redundancy, with RNF157 being able to compensate for the loss of MGRN1 in certain cell types^2^. Thus, to identify additional MGRN1 adapters, we performed co-immunoprecipitation experiments followed by mass spectrometry (co-IP/MS) using *Mgrn1^-/-^*; *Rnf157^-/-^* NIH/3T3 cells stably expressing MGRN1 fused to a 1D4 tag (MGRN1-1D4) (**Figure 1A**). This approach revealed nine proteins that interacted exclusively with MGRN1 compared to wild-type controls (**Figure 1B**). These interacting proteins included tubulin subunits (TUBB3, TUBB6), ATP synthase subunits (ATPA, ATPB), and mitochondrial proteins (CMC1, ADT2), supporting the known roles of MGRN1 in regulating microtubule dynamics^10,11^ and maintaining mitochondrial homeostasis^12–15^.

**Figure 1.**
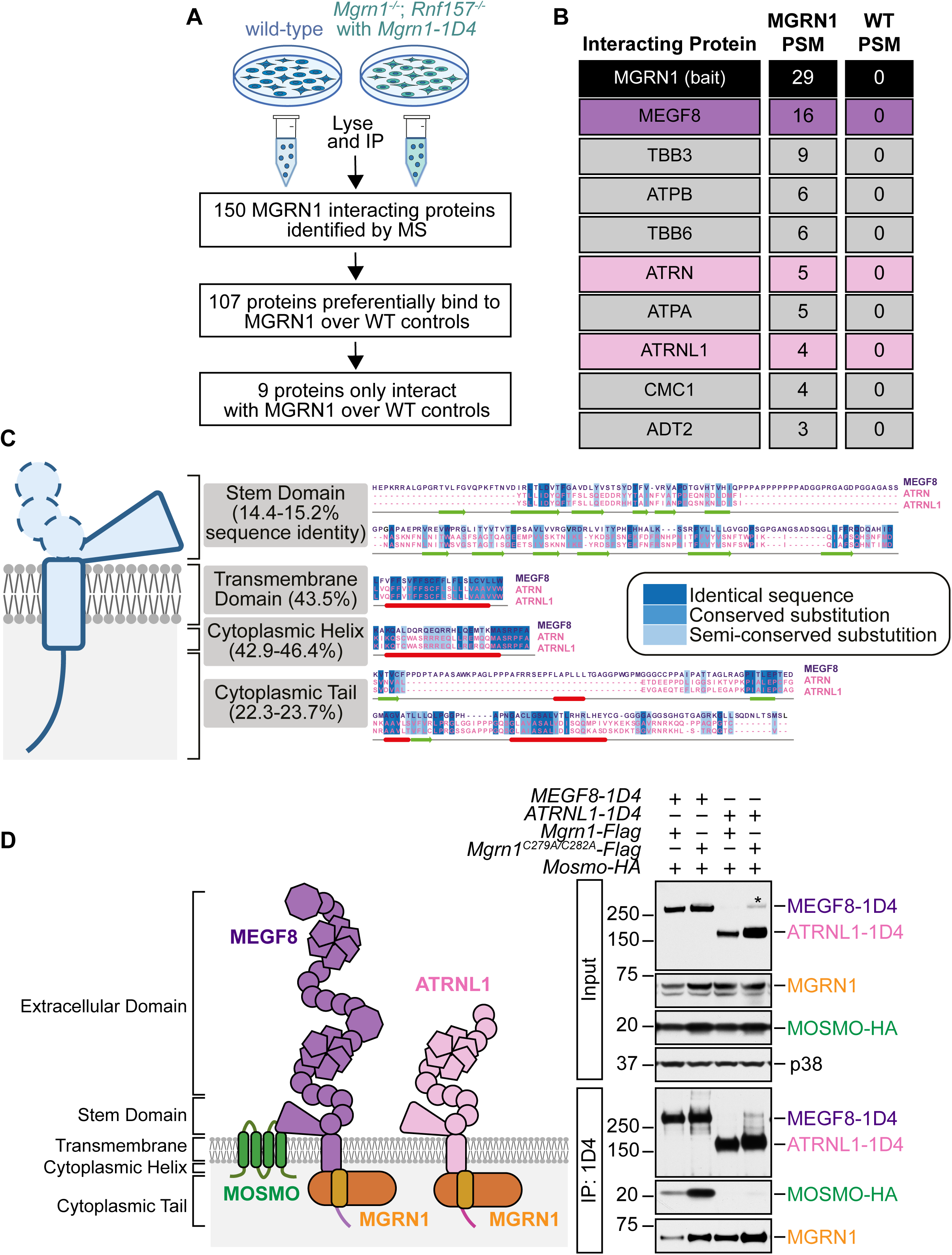
Identification of MGRN1 interacting proteins. **(A)** Schematic of the co-immunoprecipitation mass spectrometry (co-IP/MS) strategy used to identify MGRN1 interacting proteins. **(B)** Table listing nine proteins that interact exclusively with MGRN1 compared to wild-type controls. Peptide spectrum match (PSM). **(C)** Multiple sequence alignment (MSA) of the stem domain, transmembrane domain, cytoplasmic helix, and cytoplasmic tail of MEGF8, ATRN, and ATRNL1. MSA was performed using ClustalO and visualized with Jalview. Sequence identities between MEGF8 and ATRN/ATRNL1 were calculated using ClustalO. Secondary structure prediction of MEGF8 was generated with JPred, where helices are represented by red tubes and sheets by green arrows. **(D, left)** Schematic representation of the MOSMO-MEGF8-MGRN1 (MMM) and the ATRNL1-MGRN1 complexes. **(D, right)** Co-IP analysis of MEGF8 and ATRNL1 interactions. HEK293T cells were transiently transfected with 1D4-tagged MEGF8 or ATRNL1, FLAG-tagged MGRN1 (wild-type or catalytically inactive MGRN1^C279A/C282A^), and HA-tagged MOSMO. Proteins were immunoprecipitated using anti-1D4 beads and analyzed by western blot. MGRN1 interacts with both MEGF8 and ATRNL1, while MOSMO interacts specifically with only MEGF8. The asterisk (*) indicates a likely ATRNL1 dimer or oligomer that has been observed in previous studies^48^. p38 serves as a loading control.

Importantly, our analysis identified Multiple epidermal growth factor-like 8 (MEGF8), which we previously demonstrated forms a complex with MGRN1^2^, and two additional surface proteins, ATRN and ATRNL1 (**Figure 1B**).

We analyzed the sequence homology between MEGF8, ATRN, and ATRNL1 to understand the basis of their interaction with MGRN1. In humans and mice, ATRN and ATRNL1 are paralogs of MEGF8^16^. Sequence alignment revealed that most of their shared homology is localized within the transmembrane and cytoplasmic domains, with a 43.5% shared sequence identity in the transmembrane domain and 42.9-46.4% in the cytoplasmic helix (**Figure 1C**).

Notably, the highly conserved cytoplasmic motif (MASRPFA), through which MEGF8 interacts with MGRN1^2^, is also present within the cytoplasmic helix of ATRN and ATRNL1^16,17^ (**Figure 1C**). Previous studies have provided evidence supporting an interaction between ATRN and MGRN1. Fluorescently labeled ATRN and MGRN1 colocalize in HEK293T cells and interactions were observed by co-immunoprecipitation (co-IP) experiments conducted in Neuro2A neuroblastoma cells^18^. Additionally, a recent micropublication showed that the MASRPF motif is required for the interaction between *Drosophila* MGRN1 and Distracted, the fly ortholog of ATRN and ATRNL1, reinforcing the importance of this motif in mediating this interaction^19^.

Based on these reports and our co-IP/MS data, we hypothesized that MGRN1 would interact with ATRN and ATRNL1 via their intracellular domains^19^. To test this, we performed interaction assays in HEK293T cells expressing tagged versions of these proteins. Co-IP experiments confirmed that MGRN1 interacts with both MEGF8 and ATRNL1 (**Figure 1D**). Interestingly, our data revealed that ATRNL1 did not interact with MOSMO, despite its similarity to MEGF8. This lack of interaction is likely due to substantial divergence in the extracellular stem domains of MEGF8 and ATRN/ATRNL1, a region that we previously reported is critical for MOSMO binding^20^. Collectively, these findings expand upon the known membrane-bound adapters of MGRN1 and simultaneously highlight the unique interaction profiles of MEGF8 and ATRNL1.

### ATRN/ATRNL1 interact with the RING domain of MGRN1

To determine which domain of MGRN1 is responsible for interacting with ATRN and ATRNL1, we generated a series of FLAG-tagged MGRN1 truncation mutants, targeting previously described interaction and functional domains (**Table 2**). These constructs included deletions of the pre-engager domain (ΔPreEngShort and ΔPreEngLong), engager domain (ΔEng), RING domain (ΔRING), and intrinsically disordered tail (ΔTail) (**Figure 2A**). Using a Flp-In system, we stably expressed either wild-type *Mgrn1* or a truncated variant in *Mgrn1^-/-^*; *Rnf157^-/-^*NIH/3T3 cells. To assess for interactions, we then performed co-IP assays using Anti-FLAG beads to pull down MGRN1 and probed for endogenous ATRN/ATRNL1. Because the cytoplasmic tails of ATRN and ATRNL1 are highly conserved, we used an antibody generated against this region to detect both proteins (**Figure 2B**). As expected, wild-type MGRN1 robustly associated with endogenous ATRN/ATRNL1 (**Figure 2C and 2D**). However, ATRN/ATRNL1 showed reduced binding to MGRN1^ΔPreEng^, MGRN1^ΔEng^, and MGRN1^ΔRING^. Collectively, this suggests that all three domains are important for this interaction.

**Figure 2.**
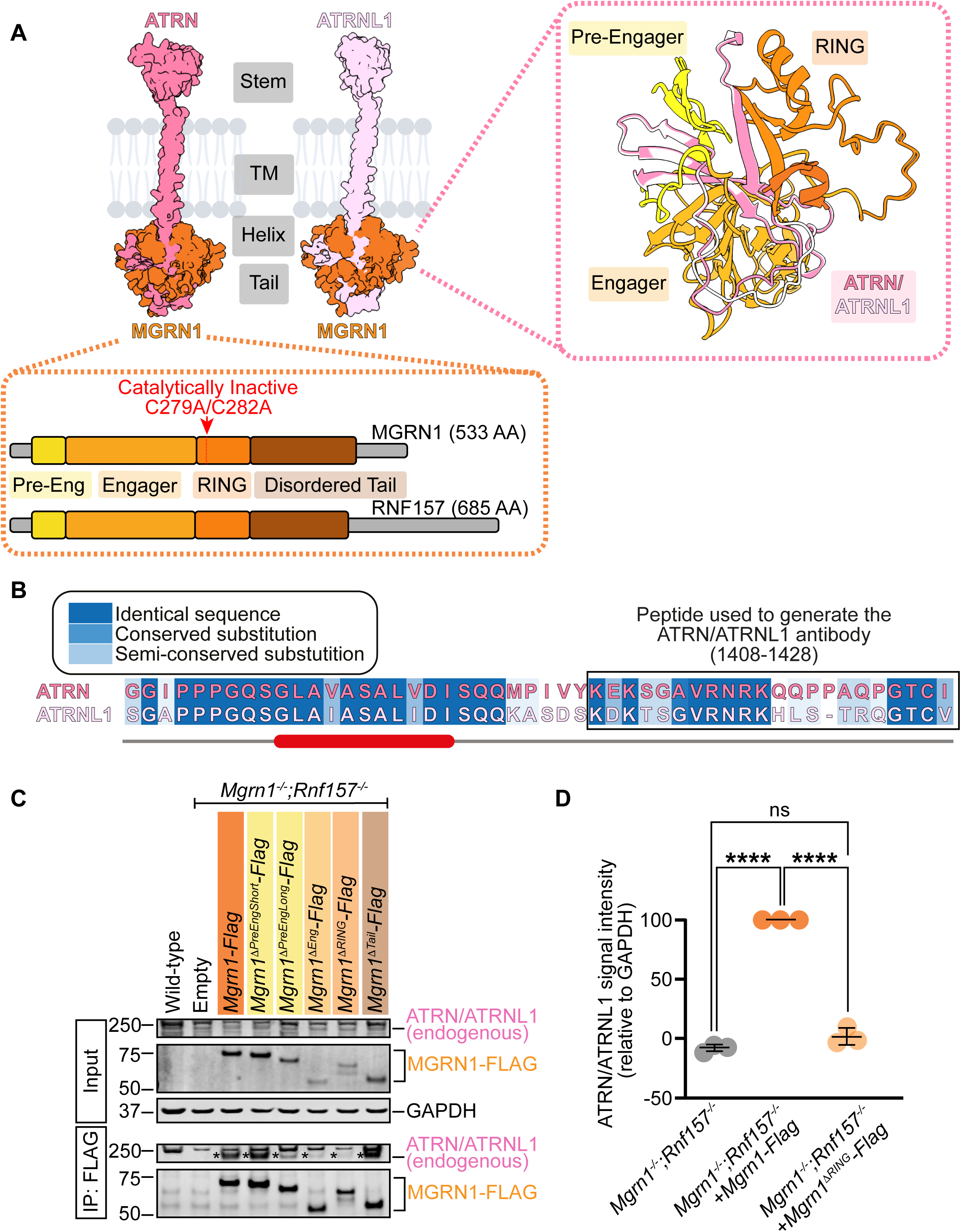
ATRN and ATRNL1 interact with the RING domain of MGRN1. **(A, Top Left)** AlphaFold-predicted ATRN-MGRN1 and ATRNL1-MGRN1 interactions. Both ATRN (residues 1157-1429, dark pink) and ATRNL1 (residues 1107-1379, light pink) contain four distinct domains, which we refer to as the extracellular stem (Stem), transmembrane (TM), cytoplasmic helix (Helix), and cytoplasmic tail (Tail). In interaction models, MGRN1 (orange) interacts with the cytoplasmic helix and tail of ATRN and ATRNL1. (**A, Bottom Left)** Schematic of the domain organization of MGRN1 (533 amino acids, 533 AA) and its vertebrate-specific paralog RNF157 (685 amino acids, 685 AA). MGRN1 and RNF157 have four previously described interaction and functional domains, which we refer to as the pre-engager, engager, RING, and intrinsically disordered tail (**Table 2**). Arrows indicate the location of the point mutations made to generate the catalytically inactive MGRN1 variant MGRN1^C279A/C282A^. **(A, Top Right)** AlphaFold prediction of ATRN and ATRNL1 interacting with the MGRN1 pre-engager, engager, and RING domains. **(B)** Multiple sequence alignment (MSA) of the cytoplasmic tail of ATRN and ATRNL1. An ATRN peptide (residues 1408-1428, boxed) that was used to generate the anti-ATRN antibody is highly conserved with ATRNL1. **(C)** Co-IP analysis of the ATRN/ATRNL1-MGRN1 interacting domains in wild-type and *Mgrn1^-/-^; Rnf157^-/-^*NIH/3T3 cells stably expressing FLAG-tagged wild-type *Mgrn1* or one of several *Mgrn1* truncation mutants. The mutants include deletions of the pre-engager *(ΔPreEngShort* and *ΔPreEngLong,* residues 35-48 and 35-82, respectively), engager (*ΔEng*, residues 79-260), RING (*ΔRING*, residues 260-335), and disordered tail (*ΔTail*, residues 335-460). Proteins were immunopreciated using anti-FLAG beads and analyzed by western blot. The expression of MGRN1 constructs in addback cell lines was verified using an anti-FLAG antibody. GAPDH serves as a loading control. The asterisks (*) indicate the endogenous monomeric band of ATRN/ATRNL1 observed in the Flag IPs. **(D)** Quantification of endogenous ATRN/ATRNL1 western blot signal intensities relative to GAPDH in immunoprecipitated samples from *Mgrn1^-/-^;Rnf157^-/-^*, *Mgrn1^-/-^;Rnf157^-/-^* with *Mgrn1-Flag*, and *Mgrn1^-/-^;Rnf157^-/-^*with *Mgrn1^ΔRING^-Flag* NIH/3T3 cell lines. The graph represents data from three independent experiments, with each dot representing one experimental replicate and with horizontal lines denoting the mean and standard deviation. Statistical significance was determined using one-way ANOVA with multiple comparisons. **** p < 0.0001 and not-significant (ns).

**Table 2:** Reported MGRN1 interactions.

| Protein | Name | MGRN1 Interacting Domain | Reference |
| --- | --- | --- | --- |
| ATRN | Attractin | Not defined | 39 |
| AMFR | Autocrine Motility Factor Receptor | Pre-engager and engager | 41 |
| HSP70 | Heat shock protein 70 | Not defined | 42 |
| MC1R | Melanocortin receptor 1 | Not defined | 6 |
| MC4R | Melanocortin receptor 4 | Not defined | 40 |
| MEGF8 | Multiple epidermal growth factor-like domains protein 8 | Not defined | 42 |
| PRIO | Major prion protein | Engager | 43 |
| TSG101 | Tumor susceptibility gene 101 protein | Disordered tail (sequence PSAP) | 44 |
| Protein | Name | CG9941 Interacting Domain (Drosophila homolog of MGRN1) | Reference |
| DSD | Distracted | Not defined | 44 |
| Protein | Name | LOG2 Interacting Domain (plant homolog of MGRN1) | Reference |
| GDU1 | Glutamine dumper 1 | Engager | 45 |

While MGRN1 has been shown to interact with multiple proteins (**Table 2**), ATRN and ATRNL1 are the first to be identified as potential RING domain-binding partners. This interaction is unexpected, as RING domains typically mediate ubiquitin transfer rather than facilitating protein-protein interactions. While these findings are consistent with our co-IP/MS results (**Figure 1B**) and are supported by AlphaFold-predicted interactions (**Figure 2A**), further studies are needed to confirm this observation.

### The ATRNL1-MGRN1 complex ubiquitinates MC1R and MC4R

We previously found that the MEGF8-MGRN1 complex attenuates Hedgehog signaling strength by ubiquitinating the GPCR SMO and facilitating its internalization and degradation^2^. Given that MGRN1 similarly interacts with ATRN and ATRNL1, we speculated that the resulting ATRN-MGRN1 or ATRNL1-MGRN1 complexes may target other surface receptors. Extensive epistasis experiments in mice have consistently shown that MGRN1 and ATRN inhibit melanocortin signaling^3–5^, specifically signaling through the GPCRs melanocortin receptor 1 and 4 (MC1R and MC4R), as evidenced by changes in coat color and satiety (**Supplementary Figure 1B**). Thus, we focused on these two receptors. As previously reported^6^, we first utilized co-IP assays in HEK293T cells to confirm that MGRN1 interacts with MC1R and MC4R (**Figure 3A and 3B**). We then established an MCR ubiquitination assay by expressing hemagglutinin (HA)-tagged ubiquitin (HA-UB) and either mVenus-tagged MC1R or MC4R in HEK293T cells, then measuring the amount of HA-UB conjugated to the MCRs under denaturing conditions.

**Figure 3.**
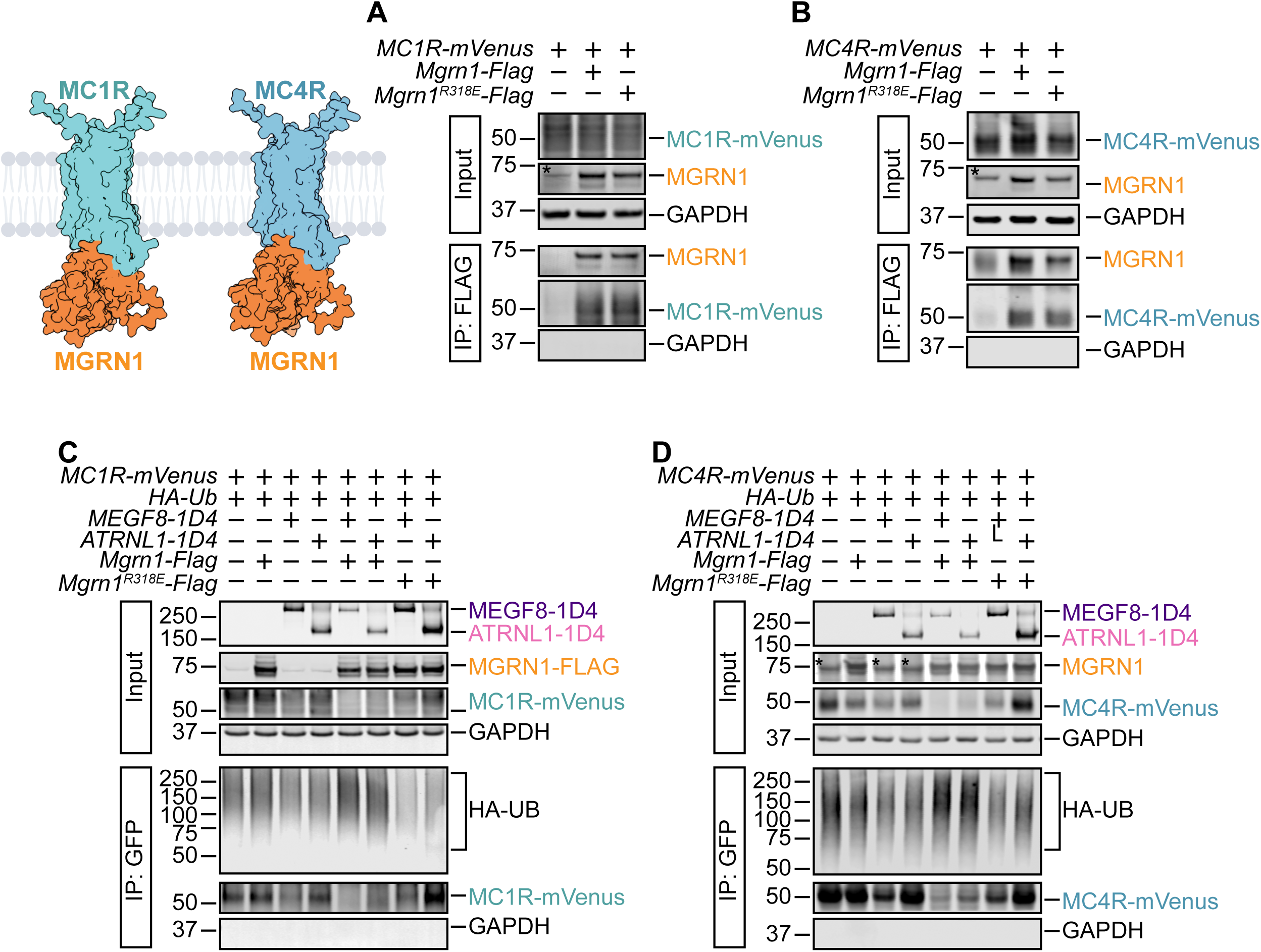
MGRN1 interacts with MC1R and MC4R and facilitates their ubiquitination. **(A, Left)** AlphaFold-predicted MC1R-MGRN1 and MC4R-MGRN1 interactions. **(A and B)** HEK293T cells were transfected with **(A)** *MC1R-Venus* or **(B)** *MC4R-Venus* along with either wild-type *Mgrn1-Flag* or the catalytically inactive *Mgrn1^R318E^-Flag* linchpin mutant. We performed co-IP assays using anti-FLAG beads to pull down MGRN1 and then probed for either MC1R or MC4R using an anti-GFP antibody. In the absence of MGRN1-FLAG, MC1R and MC4R were not detected in the pulldown. However, both wild-type MGRN1 and MGRN1^R318E^ efficiently co-immunoprecipitated with MC1R and MC4R. **(C and D)** MC1R and MC4R ubiquitination were assessed after transient co-expression of the indicated proteins in HEK293T cells. Briefly, HEK293T cells were co-transfected with **(C)** *MC1R-Venus* or **(D)** *MC4R-Venus* along with HA-tagged ubiquitin (*HA-Ub*), FLAG-tagged *Mgrn1* (wild-type or the *Mgrn1^R318E^* linchpin mutant), and either the transmembrane adapter *Megf8-1D4* or *ATRNL1-1D4*. Cells were lysed under denaturing conditions, and MC1R or MC4R was purified by IP using GFP-Trap beads. The amount of HA-UB covalently conjugated to MC1R and MC4R was assessed using immunoblotting with an anti-HA antibody. GAPDH serves as a loading control. The asterisks (*) indicate the endogenous MGRN1.

The MCRs were isolated by IP, and the attached UB chains were detected by immunoblotting with an anti-HA antibody. Co-expression of MGRN1 revealed no increase in MCR ubiquitination over baseline. Similarly, co-expression of MEGF8 or ATRNL1 alone showed no increase in MCR ubiquitination. However, the co-expression of both MEGF8 and MGRN1 or ATRNL1 and MGRN1 resulted in an increase in MCR ubiquitination and concomitantly reduced MCR abundance (**Figure 3C and 3D**). To validate these findings, we also included a catalytically inactive MGRN1^R318E^ construct. The R318E mutation disrupts the allosteric linchpin residue within the RING domain necessary for engaging ubiquitin and its E2 through hydrogen bonding (**Supplementary Figure 2A**)^21^. MGRN1^R318E^ is catalytically inactive and incapable of facilitating SMO ubiquitination. This is evident through the sustained elevation in Hedgehog signaling activity in *Mgrn1^-/-^;Rnf157^-/-^* NIH/3T3 cells with *Mgrn1^R318E^* stably added back, as measured through an elevation in ciliary SMO (**Supplementary Figure 2B** and 2C) and elevation in baseline levels of Hedgehog signaling using *Gli1* expression as a readout (**Supplementary Figure 2D**). In our assays, even when in the presence of MEGF8 or ATRNL1, catalytically inactive MGRN1^R318E^ failed to promote MCR ubiquitination and degradation (**Figure 3C and 3D**). Collectively, these results indicate that MGRN1 requires a transmembrane surface adapter (MEGF8, ATRN, or ATRNL1) to target the MCRs. Importantly, our data also shows that MCR degradation is dependent on the ubiquitin ligase activity of MGRN1.

### Loss of *Mgrn1* promotes the accumulation of MC4R at the cell surface and primary cilium

If MGRN1 regulates the ubiquitination and degradation of MC1R and MC4R (**Figure 3**), its loss should lead to the increased surface localization of these receptors (**Figure 4A**). To test this, we used NIH/3T3 cells. NIH/3T3 cells do not endogenously express *Mc4r*^22^. Thus, we first used a lentiviral system to express *MC4R-mVenus* in either wild-type or *Mgrn1^-/-^*;*Rnf157^-/-^*NIH/3T3 cells. Flow cytometry revealed that *Mgrn1^-/-^*;*Rnf157^-/-^*cells exhibited approximately 3 times higher MC4R-mVenus fluorescence than wild-type cells, suggesting that a loss of these E3 ligases results in more MC4R (**Figure 4B**). To further assess this, we transiently co-transfected *MC4R-mVenus* and *MRAP2-Flag* into wild-type and *Mgrn1^-/-^*;*Rnf157^-/-^* NIH/3T3 cells. Melanocortin receptor-associated protein 2 (MRAP2) has been shown to enhance MC4R ciliary localization in both neurons and inner medullary collecting duct 3 (IMCD3) cells^23^. Consistent with these studies, MRAP2 promoted MC4R ciliary localization in wild-type NIH/3T3 cells, where primary cilia were visualized using ARL13B staining (**Figure 4C**). Notably, *Mgrn1^-/-^*;*Rnf157^-/-^*cells exhibited an increase in MC4R ciliary localization compared to controls (**Figure 4D**). This increase in ciliary MC4R was accompanied by a similar increase in ciliary SMO (**Figure 4E**). Collectively, these findings suggest that MGRN1 regulates the surface abundance of multiple ciliary localized GPCRs, including MC4R and SMO.

**Figure 4.**
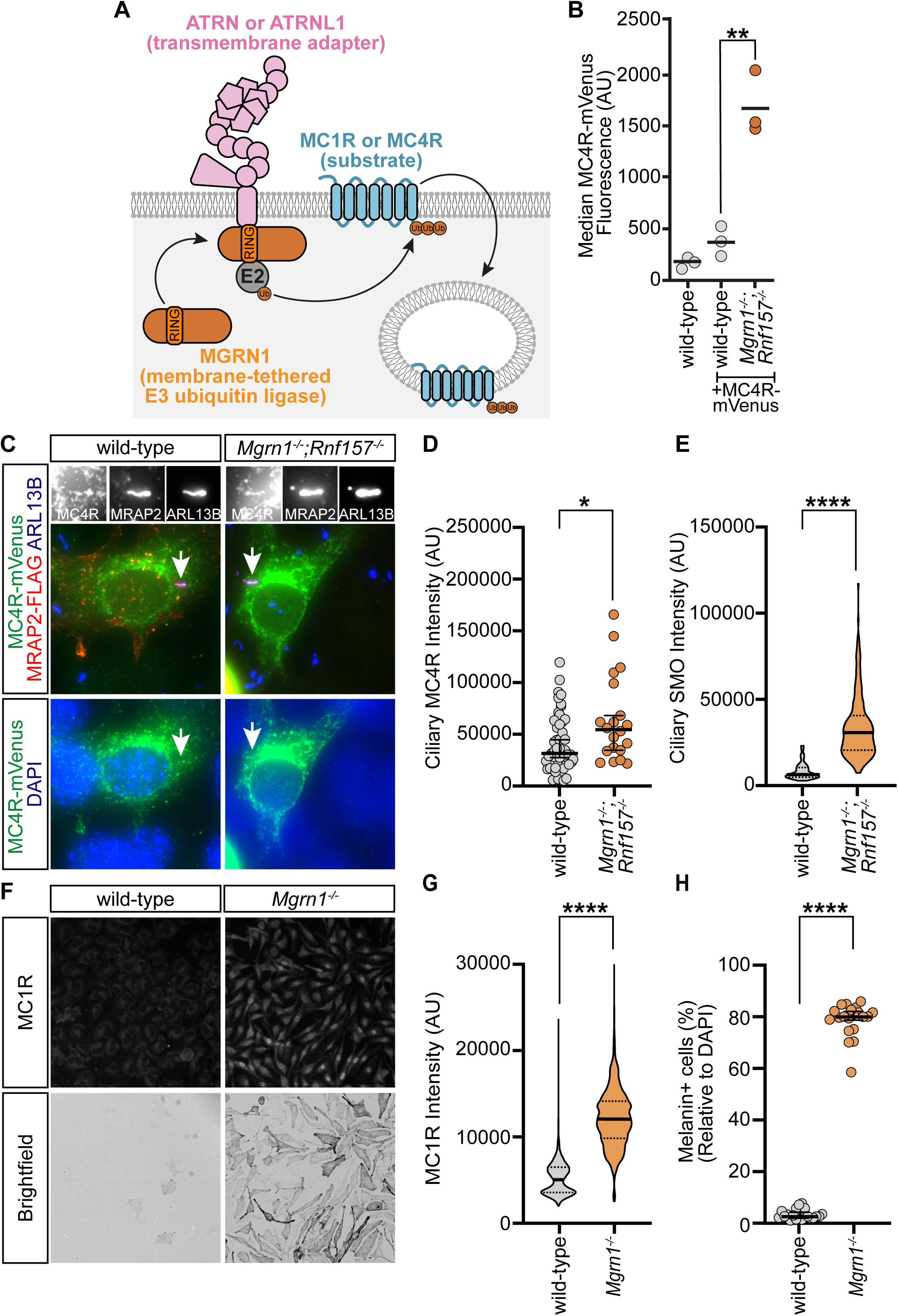
Regulation of melanocortin receptors by MGRN1. **(A)** A model depicting MGRN1 binding to the cytoplasmic tail of the transmembrane adapters ATRN or ATRNL1 to ubiquitinate the substrates MC1R or MC4R. **(B)** Flow analysis of wild-type and *Mgrn1^-/-^*;*Rnf157^-/-^*NIH/3T3 cells with stably integrated *MC4R-mVenus*. Each dot represents one experimental replicate consisting of 10,000 cells. Data is represented as a scatter dot plot and statistical significance was determined using an unpaired t-test. **p < 0.01 **(C)** Representative widefield microscopy images of wild-type and *Mgrn1^-/-^*;*Rnf157^-/-^*NIH/3T3 cells transiently transfected with *MC4R-mVenus* and *MRAP2-Flag*. ARL13B is used to identify primary cilia and DAPI to identify the nuclei. **(D)** Quantification of ciliary MC4R-mVenus in transfected wild-type and *Mgrn1^-/-^*;*Rnf157^-/-^* NIH/3T3 cells, where the primary cilia are defined by ARL13B. Each point represents single cilium. ∼20-50 cilia were analyzed per group. * p < 0.05. **(E)** Quantification of Ciliary SMO in wild-type and *Mgrn1^-/-^*;*Rnf157^-/-^* NIH/3T3 cells. The violin plot represents ∼100-200 cilia analyzed in each group. **** p < 0.0001. **(F)** Representative widefield immunofluorescence (top) and brightfield (bottom) microscopy images of primary melanocytes collected from wild-type and *Mgrn1^-/-^* mice. **(G)** Quantification of MC1R in wild-type and *Mgrn1^-/-^* melanocytes. The violin plot represents ∼1000 melanocytes analyzed in each group. **** p < 0.0001. **(H)** Quantification of the percentage of melanin+ melanocytes observed in wild-type and *Mgrn1^-/-^* samples. Each point represents a single image containing ∼300-500 melanocytes. **** p < 0.0001. **(D and H)** Data is represented as a scatter dot plot. Statistical significance was determined using the Mann-Whitney test. The bold horizontal line represents the median, with the adjacent thin lines representing a 95% confidence interval. (**E and G**) Data is represented as a truncated violin plot. Statistical significance was determined using the Mann-Whitney test. The bold horizontal line represents the median, with the adjacent dotted lines representing the first and third quartiles.

### Loss of *Mgrn1* promotes the accumulation of MC1R in melanocytes

To investigate the role of MGRN1 in regulating MC1R, we analyzed primary melanocytes derived from wild-type and *Mgrn1^-/-^* mice^24,25^. Melanocytes are specialized epidermal cells responsible for pigment production, a process that is modulated by MC1R signaling. Activation of surface MC1R initiates a signaling cascade that drives the production of eumelanin (black pigment), while the absence of MC1R signaling results in the production of pheomelanin (yellow pigment). The dark eumelanic pigmentation phenotype of *Mgrn1* null mice is consistent with increased MC1R signaling. Immunofluorescence staining of MC1R revealed a significant increase in MC1R abundance in *Mgrn1^-/-^* melanocytes compared to wild-type controls (**Figure 4F and 4G**). The loss of *Mgrn1* also led to a dramatic increase in eumelanin production, as evidenced by the number of melanin-positive cells in *Mgrn1^-/-^* melanocytes (**Figure 4F and 4H**). Collectively, this data suggests that MGRN1 plays a crucial role in modulating the abundance of MC1R, thereby influencing pigmentation outcomes.

## DISCUSSION

MGRN1 is an E3 ubiquitin ligase that was originally identified as the gene mutated in *mahoganoid* mice^16,26^. Loss of *Mgrn1* results in a broad spectrum of phenotypes, including left-right patterning defects, congenital heart malformations, embryonic lethality, craniofacial abnormalities, pigmentation defects, late-onset spongiform neurodegeneration, mitochondrial dysfunction, and partial suppression of diet-induced obesity^2–5,8,12,13,15,27–31^ (**Table 1**). While our recent work identified MGRN1 as a component of the MOSMO-MEGF8-MGRN1 (MMM) complex that regulates SMO abundance within the Hedgehog signaling pathway^2^, this function alone cannot fully explain the diverse phenotypes observed in *Mgrn1* mutant mice, suggesting that MGRN1 must regulate additional signaling pathways. To investigate this broader role, we examined MGRN1’s newly identified interactions with the transmembrane adapters ATRN and ATRNL1 (**Figures 1 and 2**). *Mgrn1* and *Atrn* null mutant mice share strikingly similar phenotypes, including a dark coat color due to the loss of a subapical band of pheomelanin on individual hairs and an elevated basal metabolic rate that confers resistance to diet-induced obesity (**Supplementary Figure 1B**, **Tables 1 and 4**). In addition, extensive epistasis experiments in mice have shown that both MGRN1 and ATRN function as negative regulators of melanocortin signaling, likely targeting MC1R and MC4R (**Supplementary Figure 1A**).

However, the precise mechanism by which MGRN1 and ATRN modulate melanocortin signaling remains unresolved. While previous studies have suggested that MGRN1 inhibits MC1R and MC4R signaling independently of ubiquitination by competing with Gαs proteins for receptor binding^6^, this model does not fully account for all the *in vivo* phenotypes observed in *Mgrn1* mutant melanocytes and mice, as previously reviewed^32^. In contrast, our findings support an adapter-mediated model of substrate specificity in which MGRN1 interacts with ATRN and ATRNL1 to regulate melanocortin receptor abundance and signaling activity. This mechanism is analogous to how MGRN1 interacts with MEGF8 to regulate SMO abundance^2,20^ and thus builds upon our understanding of the fundamental principles of membrane-tethered E3 ligases, illustrating that interactions with transmembrane adapters allow this unique group of E3s to selectively target surface receptors for ubiquitination and degradation.

One major unresolved question is how these membrane-tethered E3 complexes are regulated. In the Wnt signaling pathway, R-spondin ligands bind to the extracellular domain of the transmembrane E3 ligases ZNRF3 and RNF43 to regulate the abundance of surface receptors^33,34^. We previously speculated that the MMM complex might be regulated in a similar manner through ligands interacting with the extracellular domain of MEGF8, but to date, no such ligand has been identified. In contrast, ATRN is already known to interact with specific ligands, providing an opportunity to investigate whether these interactions influence ATRN-MGRN1-mediated degradation of melanocortin receptors. Agouti-signaling protein (ASIP) is a unique ligand because it binds to both ATRN and the melanocortin receptors MC1R and MC4R^35–37^.

Classically, ASIP is believed to function as a competitive antagonist, directly binding to MC1R and MC4R to inhibit receptor activity while also preventing receptor activation by agonists. The prevailing model is that ATRN acts as a co-receptor, facilitating ASIP’s interaction with MC1R and MC4R to inhibit signaling^37^. However, emerging evidence suggests that ASIP may play a more complex regulatory role. Although purified ASIP binds to MC1R and MC4R with high affinity in biochemical assays^38,39^, genetic studies have shown that loss of ATRN or MGRN1 disrupts ASIP-dependent MC1R and MC4R degradation, leading to the accumulation of these receptors at the cell surface^5,40^. These findings suggest that ASIP may act as an initiation factor for the ATRN-MGRN1 complex, potentially activating the complex or enhancing its ability to target MC1R and MC4R for ubiquitination and degradation. Future investigation into this model will help us understand the ligand-mediated regulation of membrane-tethered E3 ligases.

Ultimately, understanding ASIP’s role in this process will not only clarify how membrane-tethered E3s are regulated but also provide a framework for designing synthetic protein binders that modulate signaling activity by controlling receptor abundance.

An additional unresolved question is how transmembrane adapters confer substrate specificity to MGRN1. In our ubiquitination assays using overexpressed proteins, both the MEGF8-MGRN1 and ATRNL1-MGRN1 complexes were capable of targeting MC1R and MC4R. However, we suspect this interchangeability is a product of overexpression and does not reflect the biological specificity observed in tissues. The distinct, non-overlapping phenotypes of *Megf8* and *Atrn* mutant mice (**Tables 3 and 4**) strongly suggest that MEGF8 and ATRN have unique *in vivo* functions. One explanation is that differences in adapter expression levels or tissue distribution dictate substrate specificity. We see evidence of this with ATRN and ATRNL1.

**Table 3:**
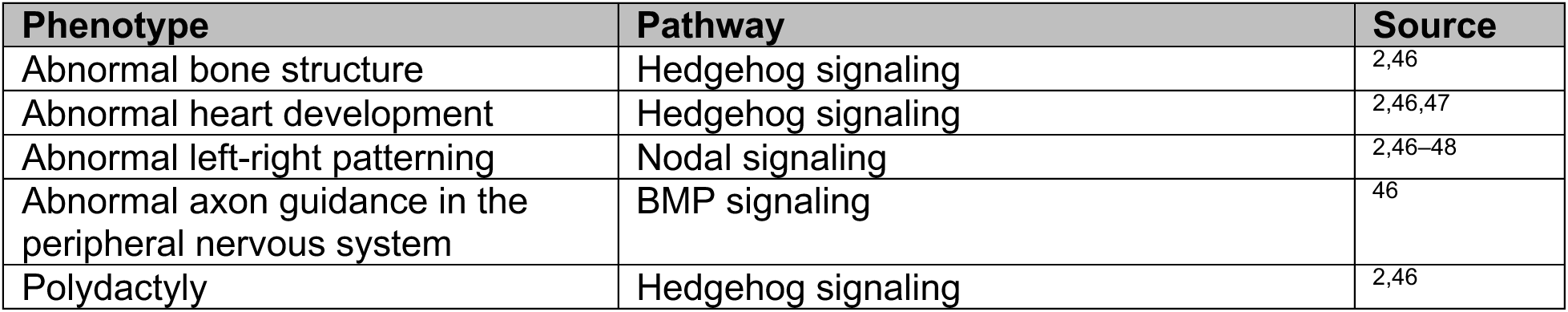
Table of phenotypes in *Megf8* mutant mice.

**Table 4:** Table of phenotypes in *Atrn* mutant mice.

| Phenotype | Pathway | Source |
| --- | --- | --- |
| Abnormal hair pigmentation | Melanocortin signaling | 49 |
| Elevated basal metabolic rate and suppresses obesity in yellow obese mouse strain (A<sup>y</sup>) | Melanocortin signaling | 3,16,37,49,50 |
| Spongiform neurodegeneration | Mitochondrial function | 40 |
| Hypomyelination | Unknown | 41 |

ATRNL1 is expressed at lower levels than ATRN, resulting in *Atrn* null mice developing pigmentation defects, an elevated metabolic rate, and neurodegeneration (**Table 4**). However, when *Atrnl1* is overexpressed under a stronger promoter (β-actin), it is able to rescue *Atrn* loss, revealing that functional redundancy can be masked by differential gene dosage^17^. Another possibility is that interaction with transmembrane adapters may change the localization of MGRN1. In this example, MEGF8 and ATRN may localize MGRN1 to distinct subcellular compartments, thus enabling selective access to specific substrates. Lastly, it is also possible that interactions with adapters may serve to activate MGRN1, ensuring that its E3 ligase activity is engaged only when appropriately localized. Such activation could involve conformational changes or post-translational modifications triggered by the adapter, acting as a safeguard to prevent off-target ubiquitination. In summary, overexpression may mask a more nuanced selection preference, highlighting the need for further exploration in endogenous systems.

Collectively, our findings provide a framework for understanding how MGRN1, as part of a membrane-tethered system, achieves substrate specificity and regulates surface receptor abundance. This work builds on the concept of adapter-mediated ubiquitination, offering a new perspective on how MGRN1 functions within signaling pathways. Further investigation into the role of transmembrane adapters will be critical for deciphering the molecular mechanisms underlying MGRN1’s diverse functions and for exploring its potential as a target to regulate aberrant signaling.

## ACKNOWLEDGEMENTS

We thank Will Walker for his helpful discussions. We also thank Rajat Rohatgi and Laura Nocka for their careful review of our manuscript. JHK was supported by the National Institutes of Health (NIH/NIGMS GM132518 and NIH/NCI CA015704) and the Dick and Anne Schneider Foundation. TMG was supported by the McLaughlin Research Institute and its generous supporters. MR was supported by the University of Washington’s Proteomics Resource (UWPR95794). MK was supported by the Howard Hughes Medical Institute (HHMI) Hanna H. Gray Fellows Program and the National Institutes of Health (NIH/NIDDK 5DP5OD03615502). CS was supported by Cancer Research UK (C20724/A26752 and DRCRPG-May23/100002). CW was supported by a DPhil studentship funded by the Wellcome Trust.

## MATERIALS AND METHODS

### Reagents and antibodies

The following primary antibodies were used: mouse monoclonal anti-1D4 (The University of British Columbia, RRID: AB_325050), guinea pig polyclonal anti-ARL13B^51^, mouse monoclonal anti-ARL13B (Aves Labs, Cat #75287, RRID: AB_2341543), rabbit polyclonal anti-ATRN (generated against a peptide from the cytoplasmic tail of ATRN, Dr. Gregory Barsh)^49^, sheep polyclonal anti-ATRN (R&D systems, Cat #AF7238, RRID: AB_2843678), chicken polyclonal anti-FLAG (Aves Labs, Cat #ET-DY100, RRID: 2313510), chicken polyclonal anti-GFP (Aves Labs, Cat #GFP-1010, RRID: AB_2307313), rabbit anti-GAPDH (LI-COR, Cat #926-42216, RRID: AB_2814901), rabbit polyclonal anti-HA (Proteintech, Cat #51064-2-AP, RRID: AB_11042321), mouse monoclonal anti-HA (Proteintech, Cat #66006-2-Ig, RRID: AB_2881490), mouse monoclonal anti-HA (Thermo Fisher Scientific, Invitrogen, Cat #26183, RRID: AB_10978021), rabbit polyclonal anti-MEGF8^51^, rabbit polyclonal anti-MC1R (Proteintech, Cat #26471-1-AP, RRID: AB_2880529), rabbit polyclonal anti-MC1R (Millipore Sigma, Cat #AB5126, RRID: AB_91692), rabbit polyclonal anti-RNF156 (anti-MGRN1, Proteintech, Cat #11285-1-AP, RRID: AB_2143351), and rabbit polyclonal anti-SMO^52^.

Secondary antibodies conjugated to IRDye® infrared dyes for western blot or Alexa Fluor dyes for immunofluorescence were purchased from LI-COR, Thermo Fisher Scientific, and Jackson Laboratories.

### Constructs

*Megf8-1D4*, *Mgrn1-Flag, Mgrn1^C279A/C282A^-Flag, Mosmo-HA* (featured in **Figures 1D and 3A-D**) were all cloned into the pEF5/FRT/V5-DEST expression vector using Gateway recombination methods (Thermo Fisher Scientific, Invitrogen) and previously described in earlier papers^52^. *pRK5-HA-Ubiquitin-WT* (referred to as “HA-Ub” in **Figures 3C and 3D**) was purchased from Addgene (Cat #17608)^53^. Full-length mouse *Mgrn1* (NM_001252437.1) with a C-terminal 3xFLAG tag was synthesized as a gBlock (Integrated DNA Technologies) and used as a template for the generation of all *Mgrn1* constructs. Overlap extension PCR was used to generate: *Mgrn1^R318E^*, *Mgrn1^ΔPreEngShort^* (deletion of aa 35-48), *Mgrn1^ΔPreEngLong^* (deletion of aa 35-82), *Mgrn1^ΔEng^* (deletion of aa 79-260), *Mgrn1^ΔRING^* (deletion of aa 260-335), and *Mgrn1^ΔTail^* (deletion of aa 355-460) (featured in **Figures 2 and 3**). Full-length Human *ATRNL1* (NM_207303.4) was PCR amplified from *pENTR223.1_ATRNL1* (Horizon Discovery) and tagged with a C-terminal 1D4 (featured in **Figures 1D**, **3C and 3D)**. All the *Mgrn1* and *ATRNL1* constructs were initially cloned into the pENTR2B plasmid (Thermo Fisher Scientific, Invitrogen) and then cloned into pEF5/FRT/V5-DEST expression vector (Thermo Fisher Scientific, Invitrogen) using Gateway recombination methods (Thermo Fisher Scientific, Invitrogen).

*MC1R-mVenus* and *MC4R-mVenus* were originally generated in the laboratory of Christian Siebold (University of Oxford, UK). Briefly, human *MC1R* (UniProt ID. Q01726) and *MC4R* (UniProt ID. P42127) genes were codon optimized and synthesized (GeneArt, Thermo Fisher Scientific). The N-terminal methionine was removed before cloning into the pHR-CMV-TetO2 vector^54^ in frame to a C-terminal 3C protease cleavage site followed by monoVenus^55,56^ and TwinStrep^57^ tags. *MRAP2-Flag* was a gift from Maxence Nachury (University of California, San Francisco). *MC1R-mVenus*, *MC4R-mVenus*, and *MRAP2-Flag* were all cloned from their respective parent plasmids into pENTR2B (Thermo Fisher Scientific, Invitrogen) and then into the pEF5/FRT/V5-DEST expression vector (Thermo Fisher Scientific, Invitrogen) using Gateway recombination methods (Thermo Fisher Scientific, Invitrogen) (featured in **Figures 3 and 4**).

### Cell Culture

The HEK293T cell line was purchased from the American Type Culture Collection (ATCC #CRL-1573) and the cells were originally isolated from the kidney of a human female embryo. The HEK293FT cell line (Cat #R70007) and the Flp-In^TM^-3T3 cell line (Cat #R76107) were both purchased from Thermo Fisher Scientific. The HEK293FT cell line is a derivative of HEK293T cells. The Flp-In^TM^-3T3 cell line is a derivative of NIH/3T3 cells, a mouse fibroblast cell line isolated from a female mouse NIH/Swiss embryo (ATCC #CRL-1658), and throughout the text we refer to these as “NIH/3T3 cells.” The HEK293T and NIH/3T3 cells were cultured in Complete Medium, composed of Dulbecco’s Modified Eagle Medium (DMEM) containing high glucose (Cytiva, Cat #SH30081.FS) and supplemented with 10% fetal bovine serum (FBS) (Thermo Fisher Scientific, Cat #A5256701), 1x GlutaMAX supplement (Gibco, Cat #35050061), 1 mM sodium pyruvate (Gibco, Cat #11360070), 1x MEM non-essential amino acids solution (Gibco, Cat #11140076), and 1x penicillin-streptomycin (Gibco, Cat #15140163). The NIH/3T3 and HEK293T cells were passaged with 0.05% Trypsin-EDTA with phenol red (Gibco, Cat #25300062).

Wild-type melan-a melanocytes were originally a gift from Dr. Dorothy Bennett. *Mgrn1^md-nc/md-nc^* (melan-md3) mutant melanocytes were originally generated by Dr. Elena Sviderskaya at the Wellcome Trust Functional Genomics Cell Bank, in collaboration with Drs. Gregory Barsh and Dorothy Bennett as described previously^25^. Immortalized melanocytes were cultured as previously reported with minor modifcations^55^. Briefly, the melanocytes were cultured on 6-well polystyrene plates coated with 0.1% gelatin (Sigma Aldrich, Cat #501785182) in RPMI 1640 basal medium (Gibco, Cat #11875093) supplemented with 1x GlutaMAX supplement (Gibco, Cat #35050061), 1x penicillin-streptomycin (Gibco, Cat #15140163),10% fetal bovine serum (FBS) (Thermo Fisher Scientific, Cat #A5256701), and 200 nM 12-*O*-tetradecanoyl phorbol-13-acetate (TPA) (Millipore Sigma, Cat #5005820001). Melanocytes were passaged upon reaching 80-85% confluency using 0.05% Trypsin-EDTA with phenol red (Gibco, Cat #25300062). All cells were housed at 37°C in a humidified cell culture incubator containing 5% CO2. All cell lines and new lines derived from these cells were free of mycoplasma contamination as determined by PCR using the Universal Mycoplasma Detection Kit (ATCC, Cat #30-1012K).

### Generation of stable NIH/3T3 cell lines expressing desired transgenes

Clonal *Mgrn1^-/-^;Rnf157^-/-^* NIH/3T3 cells were previously generated and validated^57^. Flp recombinase-mediated DNA recombination (Thermo Fisher Scientific, Invitrogen) was used to stably express *Mgrn1-1D4* (featured in **Figure 1A and 1B**), *Mgrn1-Flag*, *Mgrn1^ΔPreEngShort^-Flag*, *Mgrn1^ΔPreEngLong^-Flag*, *Mgrn1^ΔEng^-Flag*, *Mgrn1^ΔRING^-Flag*, *Mgrn1^ΔTail^-Flag* (featured in **Figure 2**), *MGRN1^R318E^-Flag* (featured **Supplementary Figure 2**), or *MC4R-mVenus* (featured in **Figure 4B**) in wild-type and *Mgrn1^-/-^;Rnf157^-/-^* NIH/3T3 cells. Briefly, wild-type and *Mgrn1^-/-^;Rnf157^-/-^* NIH/3T3 cells were transfected with 2.7 µg pOG44 Flp-recombinase expression vector (Thermo Fisher Scientific, Invitrogen) and 0.3 µg of the gene of interest (cloned into the pEF5/FRT/V5-DEST expression vector) using the X-tremeGENE™ 9 DNA Transfection Reagent (Roche Molecular Systems, Cat #6365787001) diluted in Opti-MEM^TM^ Reduced Serum Medium (Gibco, Cat #31985062). After 48 hours, the cells were passaged onto 10 cm plates and selected with Complete Medium supplemented with 200 µg/ml Hygromycin B antibiotic (Gibco, Cat #10687010). The medium containing the antibiotic was replenished every 3-4 days, and selection continued for approximately two weeks or until all cells on the control untransfected plate had died.

### Transient expression of desired constructs in NIH/3T3 cells

In a 24-well plate, 100,000 cells of either wild-type or *Mgrn1^-/-^;Rnf157^-/-^*NIH/3T3 cells were plated on glass coverslips in 500 µl of Antibiotic-Free Complete Medium (Complete Medium without the addition of 1x penicillin-streptomycin). Approximately 1 hour after plating, desired constructs (*MC4R-mVenus* and *MRAP2-Flag*) cloned into the pEF5/FRT/V5-DEST expression vector were transiently transfected into the cells using X-tremeGENE™ 9 DNA Transfection Reagent (Roche Molecular Systems, Cat #6365787001) diluted in Opti-MEM^TM^ Reduced Serum Medium (Gibco, Cat #31985062). To facilitate ciliation, after the cells were confluent, the cells were transitioned to Low Serum Medium (Complete Medium containing 0.5% FBS) overnight (featured in **Figures 4C and 4D**).

### Mass Spectrometry

Protein sequence analysis by liquid chromatography-mass spectrometry (LC-MS) was conducted at the Taplin Biological Mass Spectrometry Facility (Harvard Medical School). Briefly, excised gel bands were cut into approximately three 1 mm pieces. Gel pieces were then subjected to a modified in-gel trypsin digestion procedure^58^. Gel pieces were washed and dehydrated with acetonitrile for 10 minutes and then the acetonitrile was removed. Gel pieces were then completely dried in a speed vac. The gel pieces were then rehydrated with a 50 mM ammonium bicarbonate solution containing 12.5 ng/µl modified sequencing-grade trypsin (Promega) at 4°C. After 45 minutes, the excess trypsin solution was removed and replaced with 50 mM ammonium bicarbonate solution at a sufficient volume to cover the gel pieces. Samples were then placed in a 37°C room overnight. Peptides were later extracted by removing the ammonium bicarbonate solution, followed by one wash with a solution containing 50% acetonitrile and 1% formic acid. The extracts were then dried in a speed vac for ∼1 hour. The samples were then stored at 4°C until analysis. On the day of analysis, the samples were reconstituted in 5-10 µl of HPLC solvent A (2.5% acetonitrile, 0.1% formic acid). A nano-scale reverse-phase HPLC capillary column was created by packing 2.6 µm C18 spherical silica beads into a fused silica capillary (100 µm inner diameter x ∼30 cm length) with a flame-drawn tip^59^. After equilibrating the column each sample was loaded via a Famos auto sampler (LC Packings) onto the column. A gradient was formed and peptides were eluted with increasing concentrations of solvent B (97.5% acetonitrile, 0.1% formic acid). As peptides eluted they were subjected to electrospray ionization and then entered into a Velos Orbitrap Elite ion-trap mass spectrometer (Thermo Fisher Scientific). Peptides were detected, isolated, and fragmented to produce a tandem mass spectrum of specific fragment ions for each peptide. Peptide sequences (and hence protein identity) were determined by matching protein databases with the acquired fragmentation pattern by the software program, Sequest (Thermo Fisher Scientific)^60^. All databases include a reversed version of all the sequences and the data was filtered to between a one and two percent peptide false discovery rate.

Analysis of interacting proteins (featured in **Figure 1B**) was conducted at the University of Washington. For this process, the mass spectrometry raw data files were searched using a standardized workflow implemented in Nextflow and available at https://github.com/mriffle/nf-teirex-dda (revision:3 9f01e6119d57b34e3442fc2905ea7c63874926d). Briefly, the raw data files were converted to mzML using msconvert^61^ (ProteoWizard release: 3.0.22335), searched using comet^62^ (version 2023010), post-processed with percolator^63^ (version 3.05), and uploaded to Limelight^64^ for analysis and visualization. The comet parameters file and FASTA file used to search the data are available as part of the Limelight project (https://limelight.yeastrc.org/limelight/d/pg/project/137), associated with their respective searches.

### Protein sequence analysis

FASTA sequences of *Homo sapiens* MEGF8 (UniProt ID: Q7Z7M0), *Homo sapiens* ATRN (UnitProt ID: O75882), and *Homo sapiens* ATRNL1 (UnitProt ID: Q5VV63) were obtained from UniProtKB. Multiple sequence alignments (MSAs) were generated using Clustal Omega with default settings, employing seeded guide trees and HMM profile-profile^65^. The generated alignment was visualized using Jalview 2^66^ and color-coded to indicate sequence conservation. Secondary structure prediction of MEGF8 was performed using the JPred secondary structure prediction tool^67^ (**Figures 1C and 2A**).

### Co-immunoprecipitation and Western Blotting

HEK293T and NIH/3T3 cells were rinsed in chilled 1x phosphate-buffered saline (PBS, Gold Biotechnology, Cat #P-271-50)), scraped off the plate, and then the cells were pelleted by spinning them down at 300 x g for 5 minutes. The cell pellets were then lysed in Immunoprecipitation Lysis Buffer containing 50 mM Tris (pH 8.0) (Gold Biotechnology, Cat #T-095-100 and #T-400-500), 150 mM NaCl (Thermo Fisher Scientific, Cat #7647-14-5), 1% NP-40 (United States Biological, Cat #N3500), 1 mM DTT (Gold Biotechnology, Cat #DTT100), and 1x SIGMA*FAST*^TM^ protease inhibitor cocktail (Millipore Sigma, Cat #S8830). Cells were lysed for 1 hour on a shaker at 4°C, supernatants were clarified by centrifugation at 16,000 x g for 30 minutes, and protein concentrations were quantified using the Pierce^TM^ BCA Protein Assay Kit (Thermo Fisher Scientific, Cat #23225). 1D4-tagged MGRN1, MEGF8, and ATRNL1 (featured in **Figure 1**) were captured by a 1D4 antibody (The University of British Columbia) covalently conjugated to Dynabeads^™^ Protein A (Thermo Fisher Scientific, Invitrogen Cat #10002D).

FLAG-tagged MGRN1 (featured in **Figures 2 and 3**) was captured using anti-FLAG® M2 Magnetic Beads (Millipore Sigma, Cat #M8823). Beads were added to the protein lysates and incubated overnight at 4°C. The beads were then washed thoroughly: once with Wash Buffer A (50 mM Tris at pH 8.0, 150mM NaCl, 1% NP-40, and 1 mM DTT), once with Wash Buffer B (50 mM Tris at pH 8.0, 500 mM NaCl, 0.1% NP-40, and 1mM DTT), and finally once with Wash Buffer C (50 mM Tris at pH 8.0, 0.1% NP-40, and 1 mM DTT). The proteins were then eluted by resuspending samples in 2x Protein Sample Loading Buffer (LI-COR, Cat #928-40004) supplemented with 100 mM DTT and incubated at 37°C for 30 min. All samples were then run on NuPAGE Bis-Tris precast gels (Thermo Fisher Scientific). The resolved proteins were transferred onto an Immobilon®-FL PVDF membrane (Millipore Sigma, Cat #IPFL00010) using a wet/tank blotting system (Bio-Rad Laboratories). Membranes were imaged on a LI-COR Odyssey CLx imaging system.

For quantification of interactions between MGRN1 and endogenous ATRN/ATRNL1, the integrated intensity of the protein bands of interest (ATRN/ATRNL1) was measured using ImageStudio version 6 software (LI-COR Biosciences). Signal intensities were then normalized by dividing the intensities of immunoprecipitated proteins (ATRN/ATRNL1) by a normalized housekeeping protein (GAPDH). To compare interactions across conditions, the ratios of immunoprecipitated proteins (ATRN/ATRNL1:GAPDH) from the *Mgrn1^-/-^;Rnf157^-/-^* with wild-type *Mgrn1-Flag* addback samples were normalized to a signal intensity of 100. In each experiment, the corresponding ratios of immunoprecipitated proteins in *Mgrn1^-/-^;Rnf157^-/-^* and *Mgrn1^-/-^; Rnf157^-/-^* with *Mgrn1^ΔRING^-Flag* samples were calculated relative to the functional *Mgrn1^-/-^; Rnf157^-/-^*with *Mgrn1-Flag* NIH/3T3 samples (**Figures 2C and 2D**).

### Ubiquitination Assay

4 million HEK293T cells were plated onto a 10 cm plate. Twenty-four hours after plating, the cells were transfected using PEI Prime (Polyethylenimine Hydrochloride, 1 mg/ml, Cat #Prime-AQ100-100ML). 3 ug of each construct was transfected into the cells (at a DNA:PEI ratio of 1:3). An empty plasmid construct was used as a filler to ensure that each plate was transfected with the same amount of DNA. Forty-eight hours post-transfection, the cells were pre-treated with 10 µM Bortezomib (a proteasome inhibitor, LC Laboratories, Cat #B-1408) and 100 µM Chloroquine (a lysosome and autophagy inhibitor, Cayman, Cat #14194) for 4 hours to enrich for ubiquitinated proteins. Cells were washed twice with chilled 1x PBS (Gold Biotechnology, Cat #P-271-50) and lysed in Ubiquitination Lysis Buffer A comprised of: 50 mM Tris (pH 8.0) (Gold Biotechnology, Cat #T-095-100 and #T-400-500), 150 mM NaCl (Thermo Fisher Scientific, Cat #7647-14-5), 2% NP-40 (United States Biological, Cat #N3500), 0.25% sodium deoxycholate (Thermo Fisher Scientific, Cat #J62288), 0.1% SDS (Thermo Fisher Scientific, Cat #28312), 6M urea (Thermo Fisher Scientific, Cat #A12360), 1 mM DTT (GoldBio, Cat #DTT50), 10 µM Bortezomib, 100 µM Chloroquine, 20 mM N-Ethylmaleimide (NEM, Thermo Fisher Scientific, Cat # 23030), and 1x SIGMAFAST protease inhibitor cocktail (Millipore Sigma, Cat #S8820). Clarified supernatants were diluted ten-fold with Ubiquitination Lysis Buffer B (Ubiquitination Lysis Buffer A prepared without urea) to adjust the urea concentration to 600 mM. For these assays, we assessed ubiquitination on GFP-tagged proteins. Ubiquitinated GFP-tagged MC1R and MC4R (in **Figures 3C and 3D**) were captured using a GFP Nanobody/VHH coupled to magnetic agarose beads (ChromoTek GFP-Trap® Magnetic Agarose, Proteintech Cat #gtma-20) rotating overnight at 4°C. The beads were then washed once with Ubiquitination Wash Buffer A (Ubiquitination Lysis Buffer B + 0.5% SDS), once with Ubiquitination Wash Buffer B (Ubiquitination Wash Buffer A + 1 M NaCl), and finally once again with Ubiquitination Wash Buffer A. Proteins bound to the magnetic beads were eluted in 2x LDS sample buffer (LI-COR, Cat #928-40004) containing 30mM DTT at 37°C for 30 minutes and assayed by immunoblotting with anti-chicken-GFP (Aves Labs, Cat #GFP-1010) for GFP-tagged MC1R and MC4R and anti-mouse-HA (Proteintech, Cat #66006-2-lg) for HA-tagged ubiquitin conjugated GFP-tagged MC1R and MC4R.

### Immunofluorescence staining of cells and image quantifications

NIH/3T3 cells and primary melanocytes (in **Figures 4C and 4F**) were fixed in chilled 4% paraformaldehyde (MP Biomedicals, Cat #0215014601) diluted in 1x PBS (Gold Biotechnology, Cat #P-271-50) for 10 minutes on ice, followed by 3 rinses (5 minutes each) with 1x PBS. Cells were then incubated for 1 hour in Blocking Buffer comprised of: 1% Horse Serum (Cytiva, Cat #SH3007403) and 0.1% Triton X-100 (Thermo Fisher Scientific, Cat #BP151-100) diluted in 1x PBS. The cells were then incubated in primary antibodies overnight at 4°C and secondary antibodies for 1 hour at room temperature (all antibodies were diluted in Blocking Buffer).

Coverslips were mounted on slides using ProLong™ diamond antifade mountant with DAPI (Thermo Fisher Scientific, Invitrogen, Cat #P36962). Fluorescent images were captured on a Nikon eclipse Ti2 microscope equipped with a 20X air and 60X oil immersion objective (NA 1.42). Z-stacks (∼4µm thick stacks) were acquired with uniform acquisition settings (laser power, gain, offset, frame, and image format) within a given experiment.

For the quantification of Smoothened (SMO) at cilia, .nd2 Nikon stacked images were opened in Fiji^68^ with maximum intensity Z-projection. Ciliary masks were constructed based on the signal from ARL13B and then applied to corresponding SMO images to measure the fluorescence intensity of SMO at cilia (**Figure 4E and Supplementary Figure 2C**). For ciliary MC4R quantification, ciliary masks were manually created using Fiji in only the transfected MC4R-mVenus+ cells guided by ARL13B. The masks were then applied to the MC4R channel to measure the fluorescence intensity of MC4R at cilia (**Figures 4C and 4D**). For MC1R quantification, .nd2 Nikon stacked images were opened in the Nikon software NIS-Elements. Using Nikon’s General Analysis (GA3) tools, which allows for the customizable creation of data measurement routines, we used the DAPI image to generate a mask of each cell and then applied this to the MC1R image to measure its average pixel intensity across each cell (**Figures 4F and 4G**). Lastly, melanin-positive (melanin+) cells were manually counted by blinded individuals using the cell counter plugin in Fiji (**Figures 4F and 4H**).

### Hedgehog signaling assays in NIH/3T3 cells

For all Hedgehog signaling assays, NIH/3T3 cells were first grown to confluence in Complete Medium (containing 10% FBS), then the cells were transitioned to Low Serum Medium (Complete Medium containing 0.5% FBS) for 24 hours to encourage ciliation.

Hedgehog signaling activity was measured using real-time quantitative reverse transcription PCR (qRT-PCR) (**Supplementary Figure 2D**). Briefly, RNA was extracted from the cells using a Direct-zol^™^ RNA miniprep kit with TRI Reagent® (Zymo Research, Cat #R2053). RNA concentrations were measured on a NanoDrop^™^ spectrophotometer (Thermo Fisher Scientific) and equal amounts of RNA were used as a template for cDNA synthesis using a High-Capacity cDNA Reverse Transcription Kit (Thermo Fisher Scientific, Applied Biosystems, Cat #4368814). qRT-PCR for *mGli1* and *mGapdh* was performed on a QuantStudio 5 Real-Time PCR System (Thermo Fisher Scientific) with the following custom designed primers: *mGli1* (Fwd 5’-CCAAGCCAACTTTATGTCAGGG-3’ and Rev 5’-AGCCCGCTTCTTTGTTAATTTGA-3’) and *mGapdh* (Fwd 5’-AGTGGCAAAGTGGAGATT-3’ and Rev 5’-GTGGAGTCATACTGGAACA-3’).

*Gli1* transcript levels were calculated relative to *Gapdh* and reported as a fold change across cell lines using the comparative C_T_ method (ΔΔC_T_ method).

### Fluorescence-Activated Cell Sorting (FACS)

Wild-type and *Mgrn1^-/-^, Rnf157^-/-^* NIH/3T3 cells stably expressing *MC4R-mVenus* were generated using the Flp-In system (as described above). The cells were cultured in Complete Media until they were 80-90% confluent. At this point, the cells were detached using 0.05% trypsin-EDTA. The detached cells were filtered through a 70 µm filter to obtain a single-cell suspension. Approximately 1 million cells were collected and centrifuged at 1200 rpm for 5 minutes. The cell pellet was resuspended in PBS to remove residual culture media, and this wash step was repeated once. For fixation, the cells were treated with 4% paraformaldehyde (PFA) in PBS for 10 minutes on ice. Following fixation, the cells were washed again in PBS to remove any residual PFA and resuspended in 1 ml PBS. The expression of MC4R-mVenus was assessed using the BD FACSymphony™ A3 flow cytometer and the acquired data was analyzed with BD FlowJo™ software. Data was collected from approximately 10,000 events per replicate (**Figure 4B**).

### Quantification and statistical analysis

All data analysis and graphs were generated using GraphPad Prism 10. Violin plots were created using the ‘‘Violin Plot (truncated)’’ appearance function. In Prism 10, the frequency distribution curves of the violin plots are calculated using kernal density estimation. By using the ‘‘truncated’’ violin plot function, the frequency distributions shown are confined within the minimum to maximum values of the data set. On each violin plot, the median (central bold line) and quartiles (adjacent dotted lines, representing the first and third quartiles) are labeled (**Figures 4E and 4G, and Supplementary Figure 2C**). Dot plots were created in Prism 10 using the “Scatter dot plot” appearance function (**Figures 2D, 4B, 4D, and 4H, and Supplementary Figure 2D**). Statistical significance was determined using Prism 10, with details and p-values specified in the individual figure legends. P-values were reported using the following key: not-significant (ns) p-value > 0.05, * p-value < 0.05, ** p-value < 0.01, *** p-value < 0.001, and **** p-value < 0.0001. Additional figure details regarding the n-value and statistical test applied were reported in the individual figure legends.

---

**Supplementary Figure 1:**
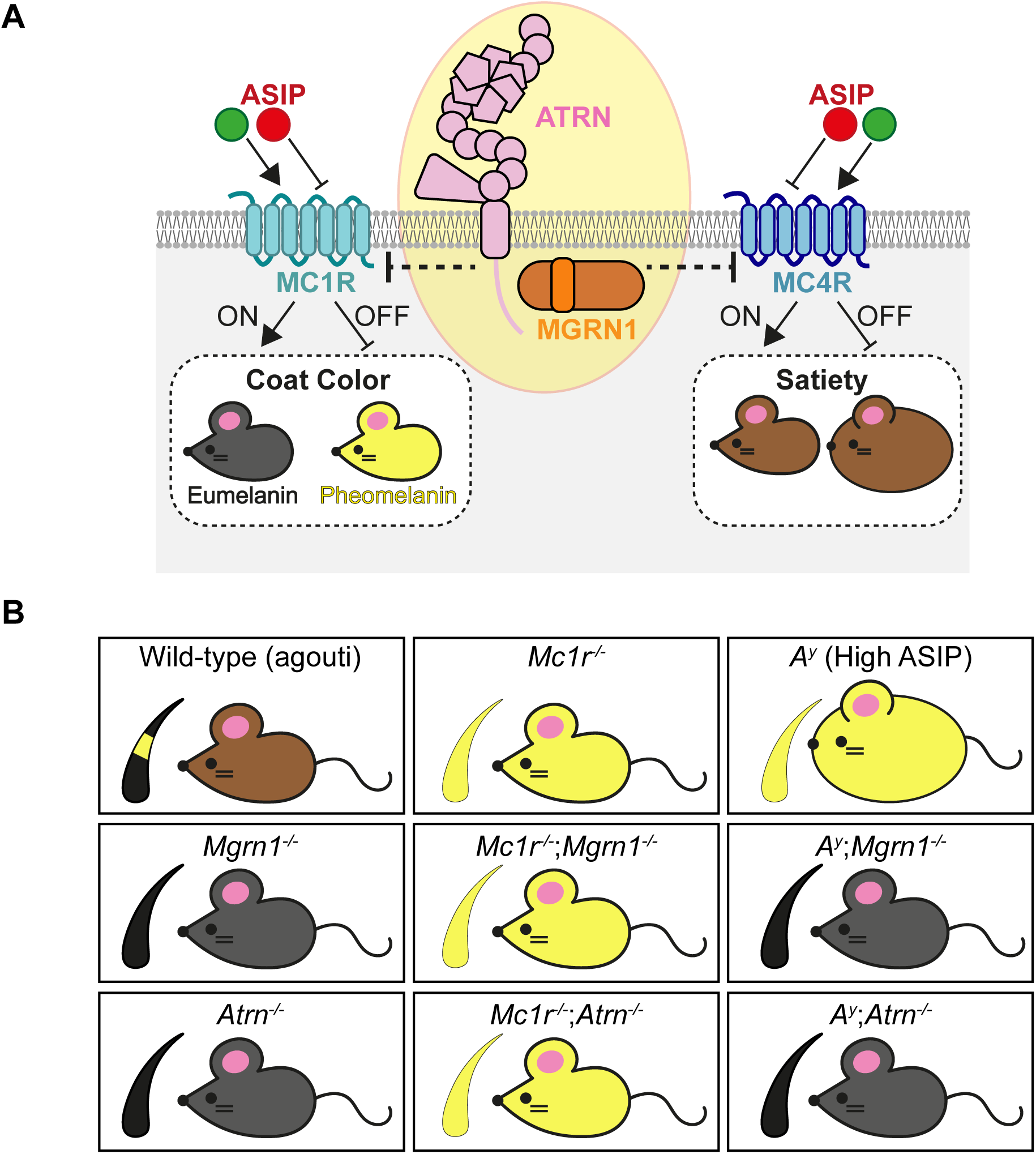
Genetic interactions show that MGRN1 and ATRN inhibit melanocortin signaling facilitated by MC1R and MC4R. **(A)** A model depicting the inhibition of melanocortin receptor function by MGRN1 and ATRN. Melanocortin receptors 1 and 4 (MC1R and MC4R) are G-protein-coupled receptors that regulate coat color and satiety. Their activity is modulated by ligands like agouti signaling protein (ASIP), which acts as an antagonist by directly binding to and suppressing receptor activity. **(B)** A summary of mouse coat color and satiety phenotypes associated with mutations in *Mgrn1, Atrn*, *Mc1r*, and *Asip*^50^. MC1R is a key regulator of coat color. When MC1R is active, it stimulates eumelanin (black pigment) production, and when inactive, leads to pheomelanin (yellow pigment) production. Wild-type (agouti) mice exhibit a banded hair phenotype, where eumelanin (black) is present at the hair base and tip, while pheomelanin (yellow) is present mid-shaft, resulting in a brown coat. Loss of function mutations in either *Mgrn1* or *Atrn* results in a darker coat due to a loss of pheomelanin production. In contrast, *Mc1r^-/-^*mutants have a fully yellow coat due to the loss of MC1R-mediated eumelanin production. The additional loss of *Mgrn1* or *Atrn* on a *Mc1r^-/-^* background has no influence on coat color, suggesting that the inhibition by MGRN1 and ATRN is at the same level or upstream of MC1R. Lastly, mice that carry the spontaneous *A^Y^* mutation overexpress ASIP, leading to inhibition of both MC1R and MC4R activity and, consequently, mice that are both yellow and obese. The additional loss of *Mgrn1* and *Atrn* on an *A^Y^* background can reverse these phenotypes, producing non-obese mice with dark coat colors. Collectively, these epistasis studies reinforce the role of MGRN1 and ATRN in suppressing melanocortin signaling via both MC1R and MC4R receptors.

**Supplementary Figure 2:**
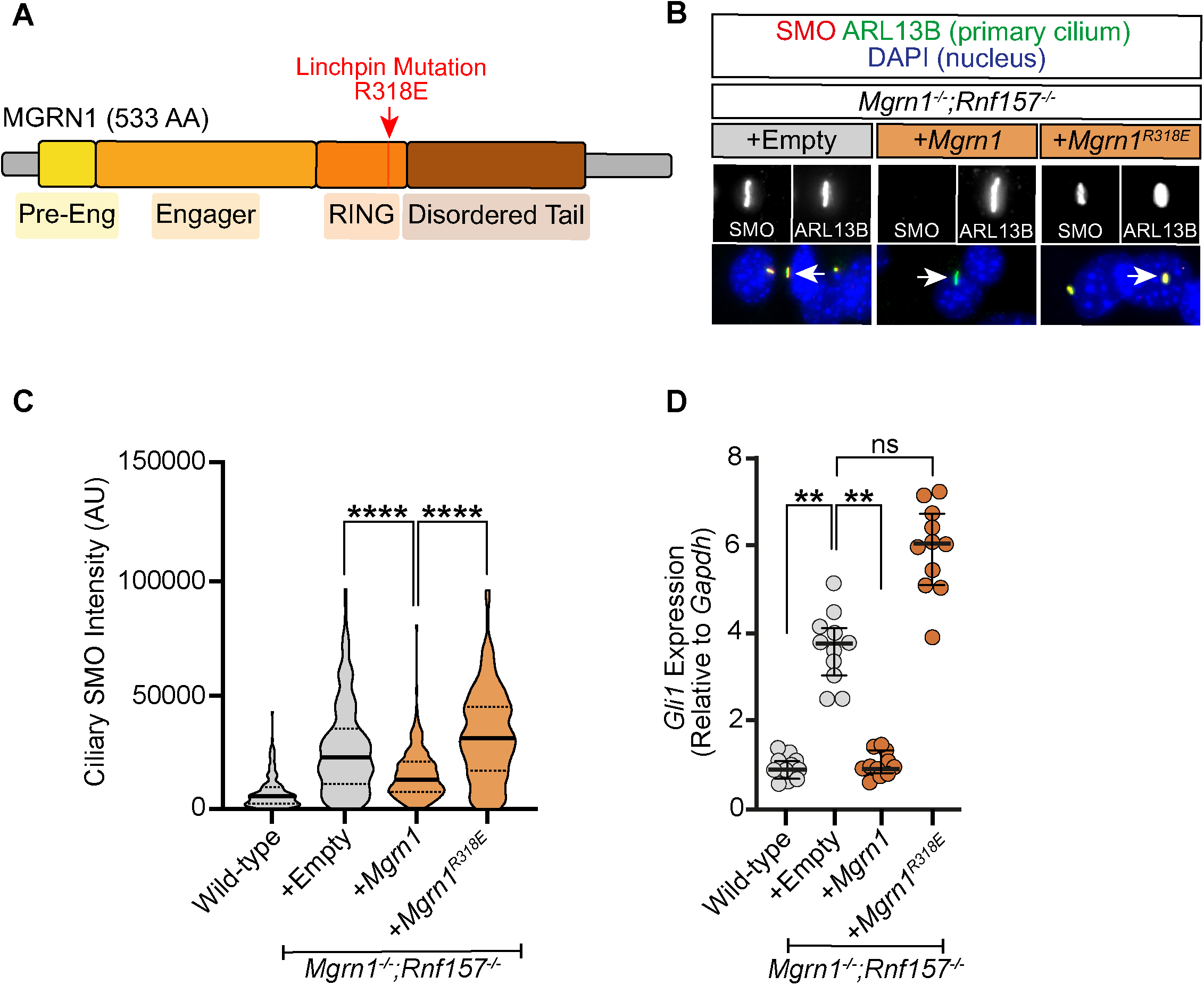
Analysis of the linchpin mutant MGRN1^R318E^. **(A)** Schematic of the domain organization of MGRN1 (533 amino acids, 533 AA). MGRN1 has four known interaction and functional domains, which we refer to as the pre-engager, engager, RING, and intrinsically disordered tail. The red arrow notes the location of the linchpin mutation (R318E) within the RING domain. **(B)** Representative widefield microscopy images of *Mgrn1^-/-^*;*Rnf157^-/-^* NIH/3T3 cells stably expressing either Flag-tagged wild-type *Mgrn1* or catalytically inactive *Mgrn1^R318E^*. SMO (red) is imaged at the primary cilia (green, marked by ARL13B), with DAPI (blue) used to label the nuclei. Arrows mark the primary cilium captured in the inset black- and-white images. **(C)** Hedgehog signaling strength was assessed by quantifying ciliary SMO in wild-type and *Mgrn1^-/-^*;*Rnf157^-/-^*NIH/3T3 cells stably expressing either wild-type *Mgrn1* or catalytically inactive *Mgrn1^R318E^*. Data is represented as a truncated violin plot of data collected from ∼200 cilia analyzed in each group. Statistical significance was determined using the Kruskal-Wallis test. **** p < 0.0001. The bold horizontal line represents the median, with the adjacent dotted lines representing the first and third quartiles. **(D)** Hedgehog signaling strength was also assessed using qRT-PCR to measure *Gli1* mRNA, a direct Hedgehog target gene. The scatter dot plot represents mRNA values derived from 3-4 individual measurements collected from three experimental replicates. The bold horizontal line represents the median, with the adjacent thin lines representing the interquartile range. ** p < 0.01 and not-significant (ns).

